# Epigenetic resetting of human pluripotency

**DOI:** 10.1101/146712

**Authors:** Ge Guo, Ferdinand von Meyenn, Maria Rostovskaya, James Clarke, Sabine Dietmann, Duncan Baker, Anna Sahakyan, Samuel Myers, Paul Bertone, Wolf Reik, Kathrin Plath, Austin Smith

## Abstract

Much attention has focussed on conversion of human pluripotent stem cells (PSC) to a more naive developmental status. Here we provide a method for resetting via transient histone deacetylase inhibition. The protocol is effective across multiple PSC lines and can proceed without karyotype change. Reset cells can be expanded without feeders with a doubling time of around 24 hours. WNT inhibition stabilises the resetting process. The transcriptome of reset cells diverges markedly from primed PSC and shares features with human inner cell mass (ICM). Reset cells activate expression of primate-specific transposable elements. DNA methylation is globally reduced to the level in the ICM but is non-random, with gain of methylation at specific loci. Methylation imprints are mostly lost, however. Reset cells can be re-primed to undergo tri-lineage differentiation and germline specification. In female reset cells, appearance of bi-allelic X-linked gene transcription indicates re-activation of the silenced X chromosome. On re-conversion to primed status, XIST-induced silencing restores monoallelic gene expression. The facile and robust conversion routine with accompanying data resources will enable widespread utilisation, interrogation, and refinement of candidate naïve cells.

## INTRODUCTION

Studies of the early mouse embryo and of derivative stem cell cultures have led to the proposition that pluripotency proceeds through at least two phases, naïve and primed (Hackett and Surani, 2014; Kalkan and Smith, 2014; Nichols and Smith, 2009; Nichols and Smith, 2012; Rossant and Tam, 2017). Recent reports provide evidence that the naïve phase of pluripotency characterised in rodent embryos may be present in a similar form in the early epiblast of primate embryos, albeit with some species-specific features (Boroviak et al., 2015; Nakamura et al., 2016; Reik and Kelsey, 2014; Roode et al., 2012; Takashima et al., 2014). However, mouse embryonic stem cells (ESC) correspond to naïve pre-implantation epiblast (Boroviak et al., 2014; Boroviak et al., 2015), while human pluripotent stem cell (hPSC) cultures (Takahashi et al., 2007; Thomson et al., 1998; Yu et al., 2007) seem to approximate primitive streak stage epiblast (Davidson et al., 2015; Irie et al., 2015; Wu et al., 2015). In general hPSC more closely resemble mouse post-implantation epiblast-derived stem cells (EpiSCs) (Brons et al., 2007; Tesar et al., 2007) than ES cells. Consequently they are considered to occupy the primed phase of pluripotency.

Mouse ESC can be propagated as highly uniform populations that exhibit consistent and unbiased multi-lineage differentiation in vitro and in chimaeras (Martello and Smith, 2014; Wray et al., 2010; Ying et al., 2008). These attributes contrast favourably with the heterogeneity and variable differentiation propensities of primed hPSC (Butcher et al., 2016; Nishizawa et al., 2016) and have provoked efforts to determine conditions that will support a human naïve condition (De Los Angeles et al., 2012). Early studies lacked stringent criteria for demonstrating a pluripotent identity with comprehensive resemblance to both rodent ESC and naïve cells in the human embryo (Davidson et al., 2015; Huang et al., 2014). However, two culture conditions have now been described for sustaining reset hPSC phenotypes that exhibit a wide range of both global and specific properties expected for naïve pluripotency (Takashima et al., 2014; Theunissen et al., 2016; Theunissen et al., 2014). Furthermore, candidate naïve hPSC can be derived directly from dissociated human ICM cells (Guo et al., 2016). These developments support the contention that the core principle of naïve pluripotency may be conserved between rodents and primates (Nakamura et al., 2016; Nichols and Smith, 2012; Smith, 2017). Nonetheless, current techniques for resetting conventional primed hPSC to a more naïve state raise issues concerning use of transgenes, universality, genetic integrity, and ease of use. Here we address these challenges and provide a simple protocol for consistent resetting to a stable and well-characterised candidate naïve phenotype.

## RESULTS

### Transient histone deacetylase inhibition resets human pluripotency

To monitor pluripotent status we exploited the piggyBac EOS-C(3+)-GFP/puro^R^ reporter (EOS) as previously described (Takashima et al., 2014). Expression of this reporter is directed by mouse regulatory elements active in undifferentiated ESC; a trimer of the CR4 element from the Oct-4 distal enhancer coupled with the early transposon (Etn) long terminal repeat promoter (Hotta et al., 2009). We observed that conventional hESC stably transfected with the piggyBac construct and maintained in KSR/FGF on feeders quickly lost visible EOS-GFP, although expression remained detectable by flow cytometry (Fig. S1A,B). Expression was further diminished when cells were transferred into 2iLIF (two inhibitors, MEK inhibitor PD and GSK3 inhibitor CH, with the cytokine leukaemia inhibitory factor, LIF; see Materials and Methods) or MEK inhibitor plus LIF (PDLIF) culture (Fig. S1C). In contrast, the PB-EOS reporter is up-regulated during transgene-induced resetting and visible expression is maintained in naïve-like cells (Takashima et al., 2014). These observations suggested that PB-EOS may be subject to reversible epigenetic silencing in primed hPSC.

Histone deacetylase (HDAC) inhibitors are global epigenetic destabilisers that have been used to facilitate nuclear transfer (Ogura et al., 2013), somatic cell reprogramming (Huangfu et al., 2008) and mouse EpiSC resetting (Ware et al., 2009). We investigated whether exposure to HDAC inhibitors would promote conversion of human primed cells to a naïve state. We applied valproic acid (VPA) or sodium butyrate to Shef6 hESC carrying the PBEOS reporter (S6EOS cells). When cells were treated for three days in E6 medium supplemented with PDLIF, then exchanged to t2iLGö naïve cell maintenance medium, the EOS reporter was up-regulated (Fig.1A,B). Bright GFP-positive colonies with dome-shaped morphology emerged over several days. We varied culture parameters and empirically determined conditions that consistently yielded per during EOS expression in compact spheroid colonies (Fig. 1A-C). We tested the method on H9EOS reporter cells and found that they similarly acquired bright GFP expression and formed dome-shaped colonies (Fig. S1D).

**Figure 1.**
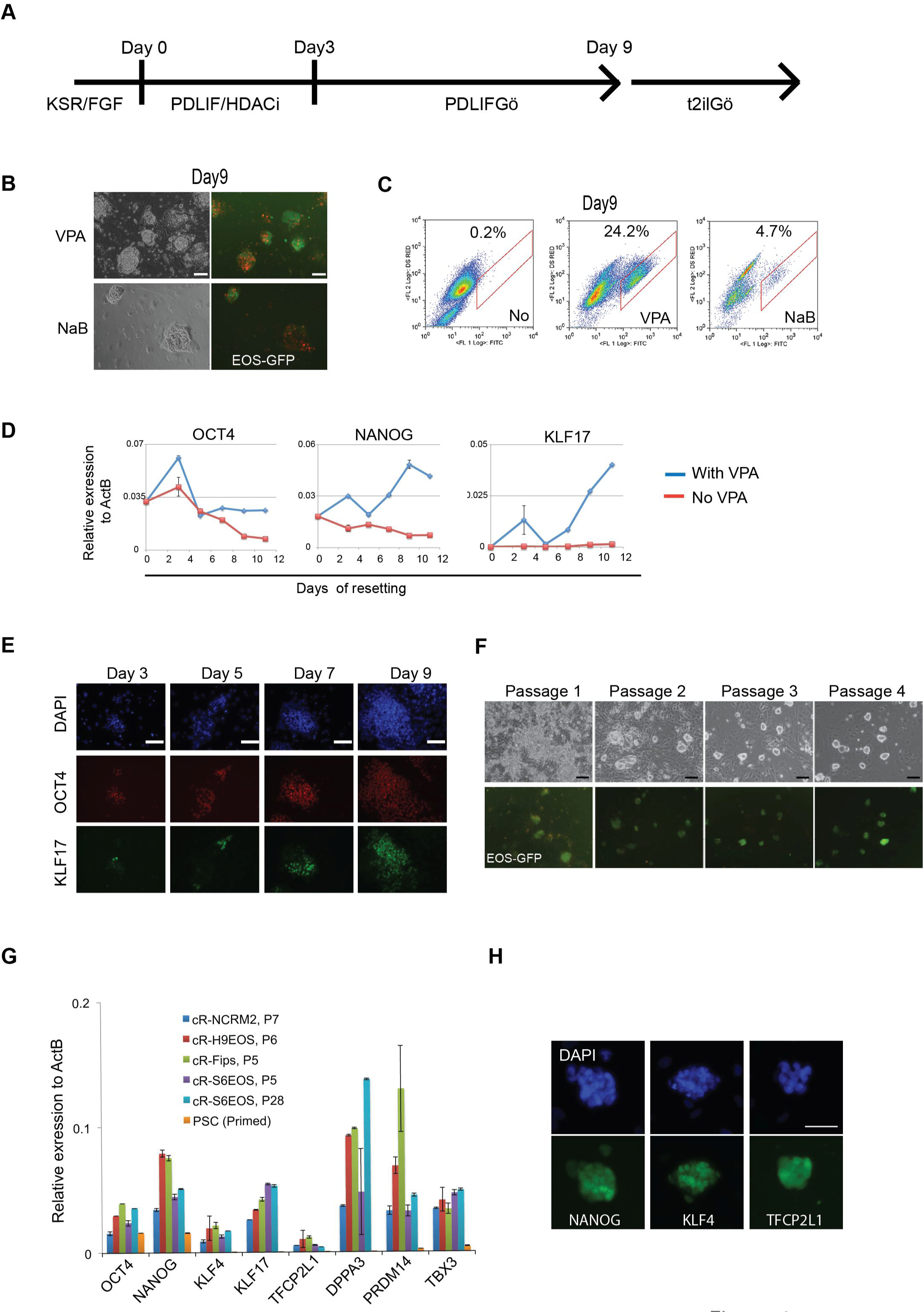
Resetting human pluripotent stem cells with HDAC inhibitors. A. Schematic of chemical resetting protocol. B. Images of reset S6EOS cells at Day 9 in t2iLGö. Red staining is from Gö6983. Scale bar, 100 μM. C. Flow cytometry analysis of EOS-GFP expression at day 9 of resetting. D. RT-qPCR analysis of pluripotency markers in S6EOS cells subjected to the resetting culture regime with or without VPA. E. Immunostaining for OCT4 and KLF17 during resetting of Shef6 cells. Scale bar, 100 μM. F. Images of reset S6EOS cultures over first 4 passages. Scale bar, 100 μM. G. RT-qPCR analysis of general and naïve pluripotency markers in various reset cell cultures. Error bars indicate SD of technical duplicates. H. Immunostaining of pluripotency markers in established reset culture, cR-H9EOS. Scale bar, 100 μM.

We monitored expression of OCT4, NANOG and the primate naïve marker KLF17 (Guo et al., 2016) during resetting of S6EOS cells. RT-qPCR analysis (Fig.1D) shows that both OCT4 and NANOG expression decrease without HDAC inhibitor treatment, consistent with differentiation in PDLIF. In contrast, in HDAC inhibitor treated cells, OCT4 mRNA levels show a transient increase on day 3 then remain at a similar level to that in primed cells, while NANOG transcripts increase about 2-fold over the first 9 days. KLF17 transcripts are not detected in conventional hESC, but become appreciable from day 7 onwards during resetting. KLF17 protein became apparent in some cells by immunofluorescence staining from as early as day 3 of resetting (Fig.1E).

Cultures were dissociated with TrypLE after nine days of resetting and replated in naïve culture media, t2iLGö. Some differentiation and cell death were evident, and a few passages were required before the EOS-positive population became stable and predominant (Fig.1F,S1E,F). From passage 5 onwards the reset phenotype was robust and could thereafter be expanded reliably.

Ability to enrich the naïve phenotype after resetting by bulk passaging in t2iLGö, suggested that a reporter should be dispensable, facilitating general applicability. We therefore tested resetting without the EOS transgene on a panel of human primed PSC. Stable cultures of compact colonies displaying naive marker gene expression were established consistently (Table 1, Fig.1G). These cell lines are denoted by the designation cR (chemically reset). Resetting efficiency varied between lines and according to initial culture status. In general, however, a single 6-well of primed PSC was sufficient for initial generation of multiple colonies and subsequent stable naïve cultures by passage 5. Rho-associated kinase (ROCK) inhibitor was used during resetting and initial expansion in most experiments but was usually omitted during subsequent expansion. Together with NANOG, reset cells expressed naïve transcription factor proteins KLF4 and TFCP2L1 that are present in the human ICM (Takashima et al., 2014) but undetectable in primed PSC (Fig.1H).

**Table 1.**
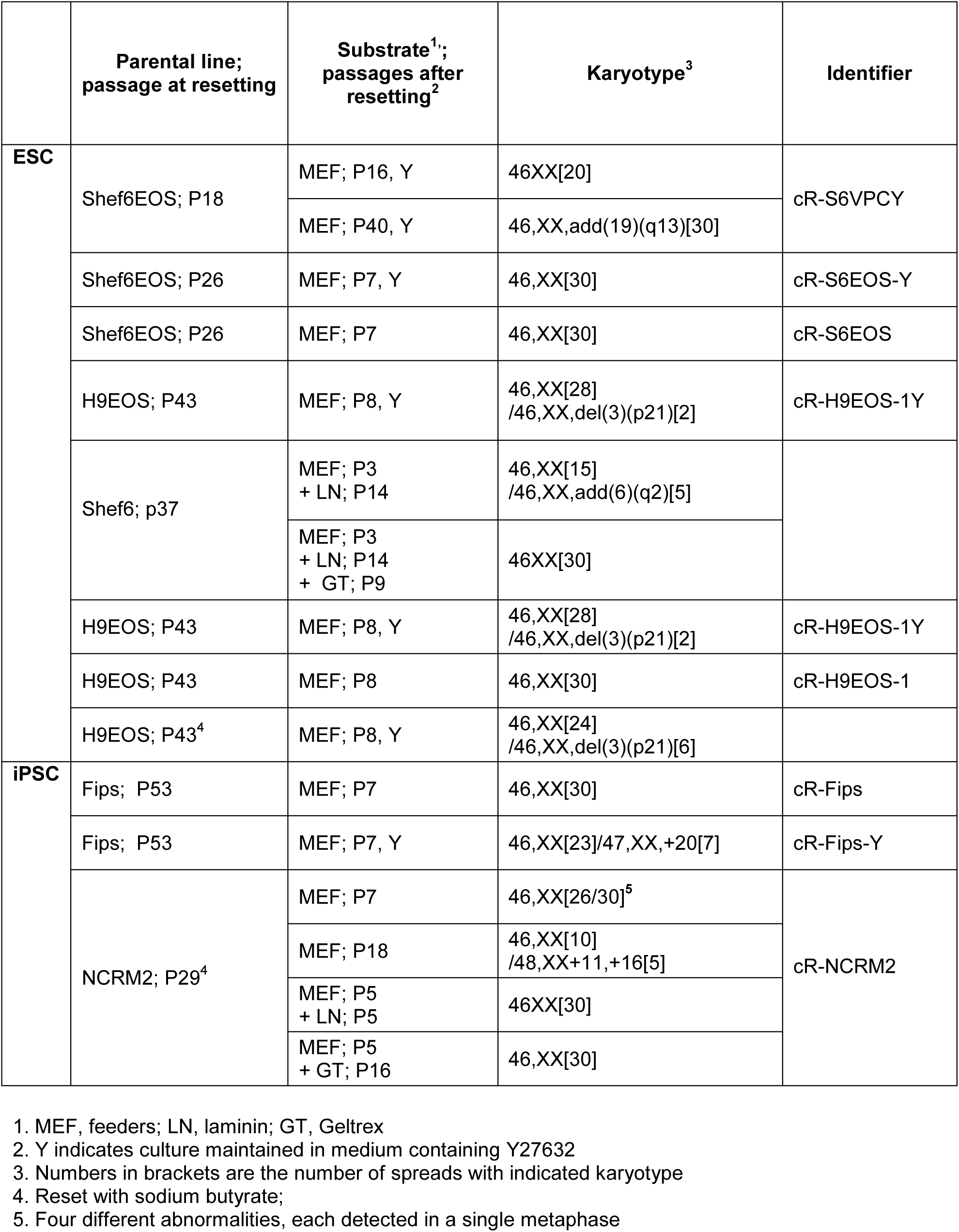
Karyotype analyses of reset cultures

### Feeder-free expansion of reset cells

As noted previously (Takashima et al., 2014), reset cells can be cultured on pre-coated plates without feeders. However, morphology was heterogeneous with more differentiation and cell death than on feeders. We varied conditions and found that provision of growth factor-reduced Geltrex with the culture medium at the time of plating was more effective than pre-coating (Fig. 2A). Geltrex or laminin applied in this manner supported continuous propagation in t2iLGö of both embryo-derived HNES and chemically reset cells with robust expression of naïve pluripotency factors (Fig. 2B-D). Moreover, aberrant expression of some mesoendodermal genes was reduced in feeder-free conditions (Fig. 2E).

**Figure 2.**
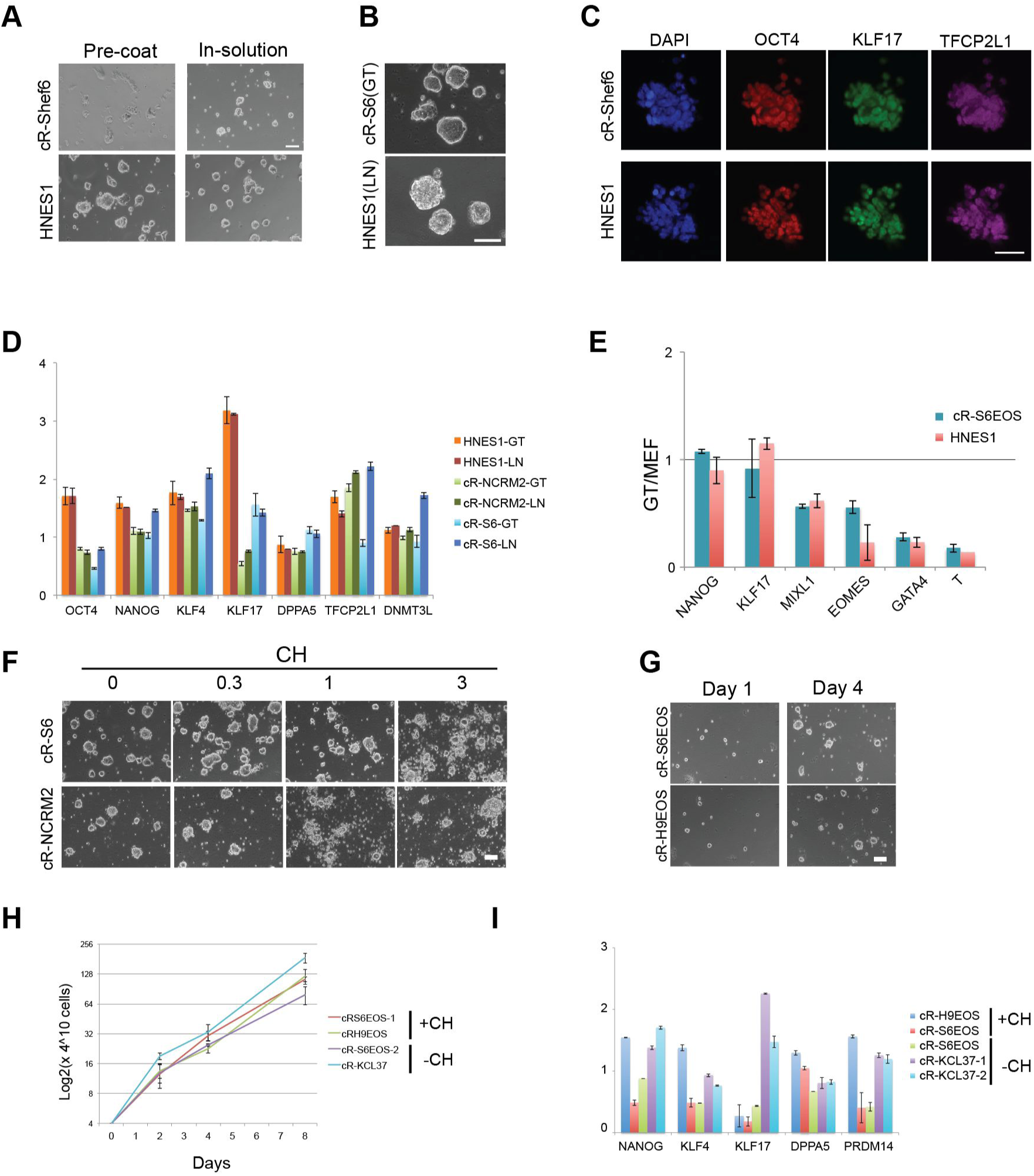
Feeder-free culture. A. Cells plated on Geltrex-coated plates (Left panels) or with Geltrex added to the medium (Right panels). Images taken after 4 days. Scale bar, 100 μM. B. Cultures in Geltrex (GT) or laminin (LN) for more than 10 passages. Scale bar, 100 μM. C. Immunostaining for pluripotency markers in reset cells passaged in laminin. Scale bar, 50 μM D. Naive marker expression in feeder-free reset cultures in t2iLGö determined by RT-qPCR and normalized to expression level in H9NK2 transgene reset cells. E. Lineage marker expression in feeder-free reset cultures relative to levels on feeders. F. Reset cells plated in the presence of indicated concentrations of CH for 4 days. Scale bar, 100 μM. G. Images of colony expansion over 4 days in Geltrex (GT). H. Growth curve for reset cells in tt2iLGö and Geltrex. I. RT-qPCR marker profile for cells reset with or without CH and expanded in tt2iLGö and Geltrex. Error bars on PCR plots indicate SD of technical duplicates.

In the absence of feeders we found that some reset cell lines expanded more robustly in very low (0.3μM) or even no CH (Fig. 2F). This is in line with observations that GSK3 inhibition is optional in the alternative 5i/L/A naïve culture system (Theunissen et al., 2016). We subsequently adopted 0.3μM CH for standard culture. Naïve cell maintenance medium with 0.3μM CH is termed tt2iLGö. Reset cultures in Geltrex and tt2iLGö displayed homogeneous morphology and expanded continuously with a doubling rate of approximately 24 hours (Fig. 2G, H).

We also observed that omitting CH entirely for the first 10 days of resetting increased the yield of EOS positive cells. We therefore implemented a revised resetting routine, omitting CH initially then exchanging into tt2iLGö on feeders before transfer to Geltrex culture. PSC reset in these conditions showed consistent feeder-free expansion with typical naïve morphology, growth and marker profiles indistinguishable from cells reset in presence of CH (Fig. 2I).

### WNT inhibition stabilises resetting

As noted above, EOS-GFP and KLF17 immunopositive colonies emerged within 10 days of VPA treatment (Fig. 1E). However, differentiation and cell death are ongoing for several passages and during this period we observed that the reset phenotype could not be sustained without feeders. Thus the resetting process appears incomplete and vulnerable at early stages. We also noted requirement for a stabilisation period following DOX-withdrawal during transgene-mediated resetting (Takashima et al., 2014). We used H9-NK2 cells, with DOX-dependent expression of *NANOG* and *KLF2,* to explore conditions that might stabilise resetting. We tested two candidates, the amino acid L-proline and the tankyrase inhibitor XAV939 (XAV). L-proline is reported to be produced by feeders and to alleviate nutrient stress in mouse embryonic stem cells (D’Aniello et al., 2015). XAV inhibits canonical Wnt signalling (Huang et al., 2009) and has previously been reported to facilitate propagation of pluripotent cells in alternative states (Kim et al., 2013; Zimmerlin et al., 2016). We withdrew DOX from H9-NK2 cells and applied either L-proline (1 mM) or XAV (2μM) in combination with t2iLGö. We assessed colony formation on feeders after the first and second passages. We saw no pronounced effect of L-proline. In contrast, addition of XAV resulted in more robust production of uniform domed colonies (Fig. 3A). RT-qPCR analysis substantiated presence of naïve pluripotency markers in XAV-supplemented cultures and also highlighted reduced levels of lineage-affiliated markers such as BRACHYURY and GATA factors (Fig. 3B).

**Figure 3.**
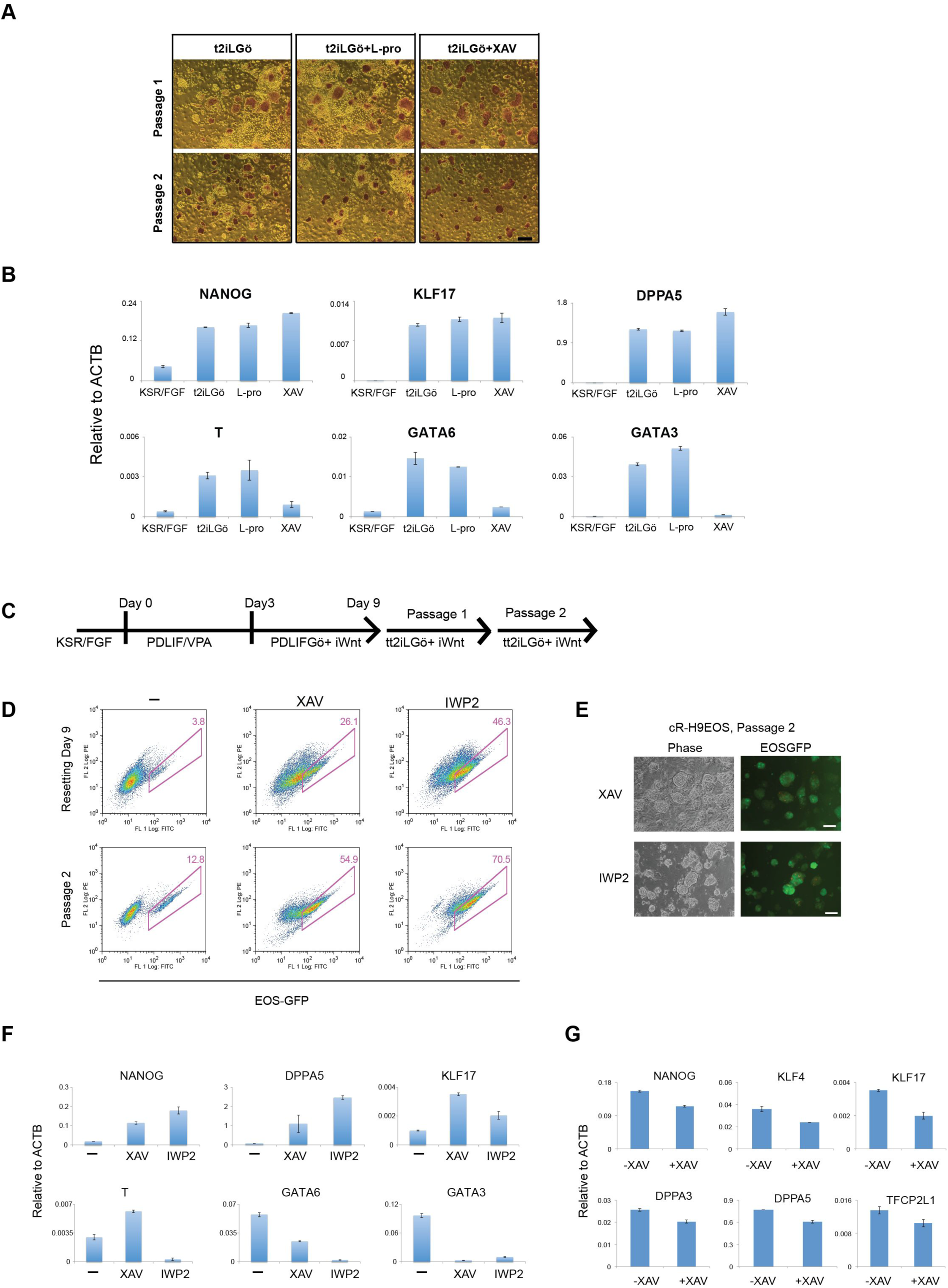
WNT inhibition stabilises resetting. A. Alkaline phosphatase staining of H9NK2 colonies at first and second passage after DOX withdrawal and transfer into t2iLGö alone or plus L-Proline (L-Pro) or XAV. Scale bar, 200 μM. B. RT-qPCR analysis of marker expression in H9NK2 cells at passage 2, treated as in A. KSR/FGF reference sample is a conventional S6EOS culture. C. Resetting protocol with WNT inhibitors. D. Top, flow analysis of resetting H9-EOS cells cultured in the presence or absence of WNT inhibitors. Bottom, flow analysis after two passages (a further 8 days) in tt2iLGö with WNT inhibitors on MEF. E. cR-H9EOS colonies in tt2iLGö with XAV or IWP2 after two passages on MEF. Scale bar, 100 μM. F. Marker analysis by RT-qPCR for cR-H9EOS cells at passage 2 cultured in tt2iLGö with and without WNT inhibitors. G. Marker analysis by RT-qPCR of cR-H9EOS cultures generated with or without XAV and transferred into tt2iLGö on Geltrex (without XAV) for 4 passages. Error bars on PCR plots indicate SD of technical duplicates.

We investigated whether WNT inhibition would stabilise emergent chemically reset cells. In addition to the tankyrase inhibitor XAV we tested an orthogonal WNT pathway inhibitor, IWP2, which acts to prevent production of functional WNT protein (Chen et al., 2009). XAV or IWP2 were added following VPA treatment on day 3 of resetting H9EOS and S6EOS cells (Fig. 3C). For both inhibitors we observed reduced numbers of differentiating or dying cells and a substantial increase in the frequency of EOSGFP positive cells by day 9 that increased further on passaging into tt2iLGö on MEF (Fig. 3D; S2A). After the second passage the majority of colonies displayed domed morphology and readily visible GFP (Fig. 3E). WNT inhibitor treated H9EOS cultures at passage 2 expressed higher levels of naïve markers and lower *GATA6* and *GATA3* than parallel cultures reset without WNT inhibition. Similarly, S6EOS cells reset using XAV or IWP2 progressed to stable reset cultures expressing naïve markers and minimal levels of BRACHYURY, CDX2 and GATA6 (Fig. S2B). From passage 3, we transferred XAV-treated cells to feeder-free culture in tt2iLGö and Geltrex without XAV. Marker analysis by RT-qPCR confirmed maintained expression of signature naïve pluripotency factors after 4 passages at similar levels to reset cells generated without use of WNT inhibitors (Fig. 3G).

We also assessed whether vitamin C was required for resetting. For the 3 day period of exposure to VPA we replaced E6 medium, which contains vitamin C, with N2B27 medium with or without addition of vitamin C. Resetting was continued in the presence of XAV as above. After two passages we observed comparable up-regulation of EOS-GFP and similar expression of naïve markers with or without exposure to vitamin C (Fig. S2C,D).

Collectively these findings establish that following VPA treatment WNT inhibition can improve the rate and efficiency of conversion to a stable naïve phenotype that can subsequently be propagated robustly in tt2iLGö with or without feeders or ongoing WNT inhibition. The results also indicate that vitamin C supplementation is not required for resetting. Full details of the protocol are provided in Supplemental Information.

### Global transcriptome profiling

We obtained transcriptome data by RNA sequencing (RNA-seq) of replicate samples of reset cells generated by VPA treatment. We also sequenced the embryo-derived naïve stem cell line HNES1 (Guo et al., 2016) and a parallel culture of HNES1 cells that had been “primed” by transfer into KSR/FGF for more than 10 passages. We added to the analysis published data (see Methods) from cells reset with inducible transgenes (Takashima et al., 2014), HNES cells cultured in the presence of vitamin C and ROCK inhibitor (Guo et al., 2016), naïve-like cells in 5i/L/A (Ji et al., 2016) and a variety of conventional PSC from publicly available resources and our own studies. We applied two complementary dimensionality reduction techniques: principal component analysis (PCA) identifies and ranks contributions of maximum variation in the underlying dataset, whereas t-distributed stochastic neighbour embedding (t-SNE) is a probabilistic method that minimises the divergence between pairwise similarities in the constituent data points. Both analyses of global transcriptomes unambiguously discriminate naïve/reset samples from primed PSC (Fig. 4A, B). In each analysis, cR cells cluster closely together with HNES1 cells that were cultured in parallel. Sample replicates are intermingled despite being from cell lines of disparate provenance and culture history. Feeder-free cultures form a slightly distinct cluster within the naïve grouping. Consistent with previous analyses (Huang et al., 2014; Irie et al., 2015; Nakamura et al., 2016; Takashima et al., 2014; Theunissen et al., 2016), two independent RNA-seq datasets for purported naïve cells cultured in 4i (NHSM) conditions (Gafni et al., 2013; Irie et al., 2015; Sperber et al., 2015) cluster with conventional primed PSC by both PCA and t-SNE, as do cultures in “extended pluripotency” media (Yang et al., 2017). For both naïve and primed cells, PCA component 2 appears sensitive to differences in growth conditions and/or batch effects and to capture variation between laboratories and cell lines.

**Figure 4.**
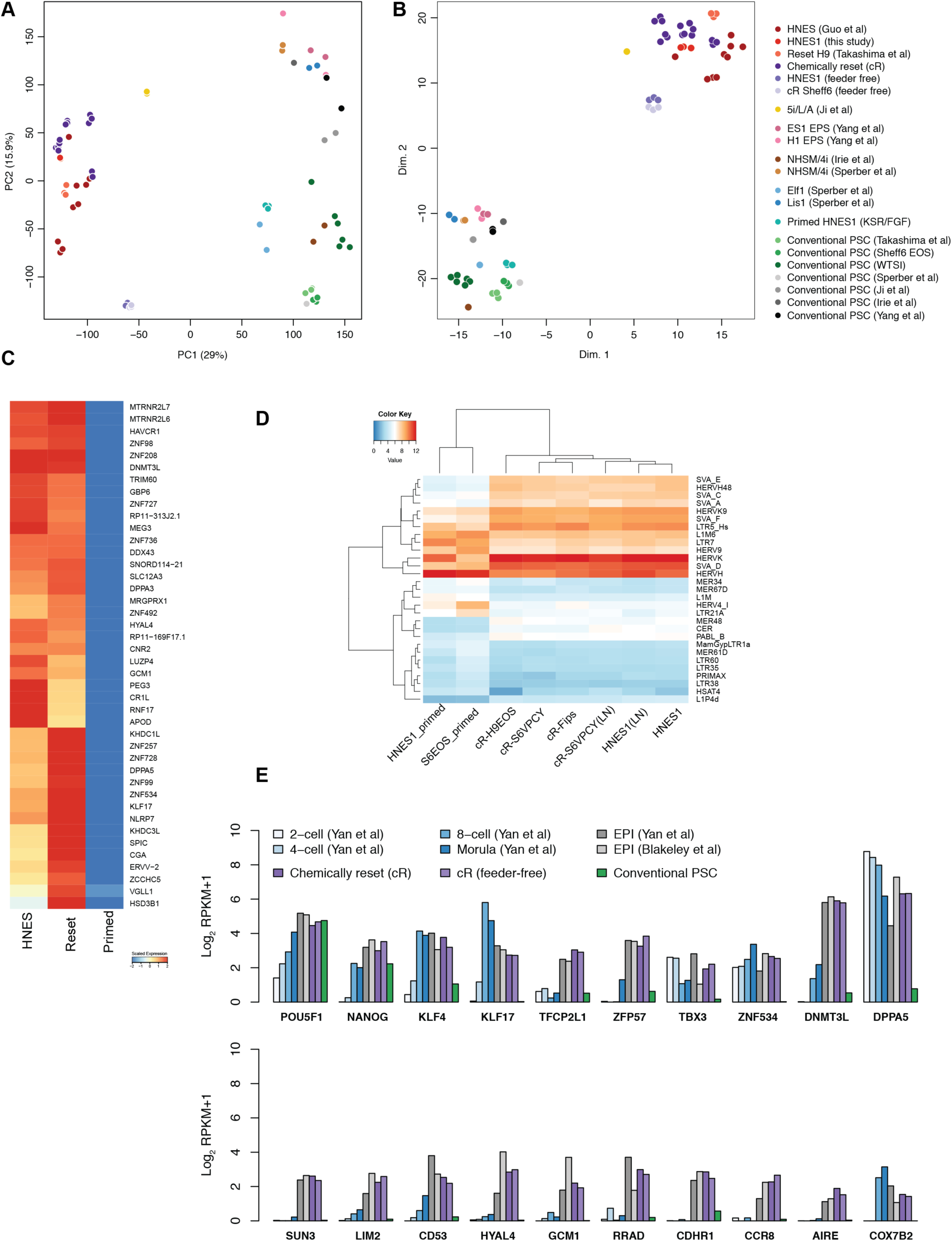
Transcriptome analysis. A. Principal component analysis of whole transcriptome RNA-seq data from indicated cell lines. B. t-SNE analysis of RNA-seq data. C. Heatmap of differentially expressed genes between chemically reset and embryo-derived HNES cells (naive) compared to conventional hPSC (primed). Genes unregulated in naive cells are shown, ranked by log2 fold-change. Values displayed correspond to the average expression level in each sample group scaled by the mean expression of each gene. D. Principal component analysis of transposon expression in naive and primed PSC. Read counts for individual TE loci were pooled for each transposon family; only TE loci with at least 10 counts were considered. E. Heatmap showing expression of all transposon families that are differentially expressed (log2FC > 1.5, P value < 0.05). F. Comparative expression of pluripotency markers in human embryo cells (Blakeley et al., 2015; Yan et al., 2013), HNES cells, cR cells, conventional primed PSC, NHSM cultures and purported expanded potency (EP) cells. Data shown reflect mean expression levels from cell lines and biological replicates belonging to each sample group, and single cells from indicated embryo stages. Published datasets used are identified in Materials and Methods.

Gene Ontology (GO) analysis of genes contributing to PC1 shows significant enrichment of functional categories primarily associated with development and differentiation (Table S2), reflecting distinct identities associated with naïve and primed cells. We also noted up-regulation of multiple genes associated with mitochondria and oxidative phosphorylation in reset cells cultured on laminin and on feeders (Fig. S3A-C) consistent with metabolic reprogramming between primed and naïve pluripotency (Takashima et al., 2014; Zhou et al., 2012). Overall cR cells share global gene expression features with ICM-derived HNES cells and transgene-reset PSC and are distinct from various primed PSC. Genes highly up-regulated in naïve conditions relative to conventional PSC are highlighted in Figure 4C.

We inspected expression of transposable elements (TEs), the transposcriptome (Friedli and Trono, 2015). A number of TEs are known to be transcriptionally active in early embryos and pluripotent stem cells, potentially with functional significance. Principal component analysis of TE expression separated cR and HNES cells from primed PSC (Fig. S3D,E). Notably, HERVK, SINE-VNTR-Alu (SVA) and LTR5_Hs elements were up-regulated in naïve cultures (Fig. 4D). Inspection of KRAB-ZNFs, potential regulators of TE expression, revealed that many are significantly up-regulated in reset cells (Fig. S3F). These include ZNF229 and ZNF534, which represses HERVH elements (Theunissen et al., 2016), ZNF98 and ZNF99, which are also up-regulated during epigenetic resetting of germ cells (Tang et al., 2015), and ZFP57, which protects imprints in the mouse (Quenneville et al., 2011).

We compared relative transcript levels for a panel of pluripotency markers between cR cells and human pre-implantation embryos. For the embryo data we used published single cell RNA-seq (Blakeley et al., 2015; Yan et al., 2013). Normalised expression was consistent between reset cells and the epiblast, more so than with earlier stage embryonic cells (Fig. 4E). Primed PSC exhibited no or low expression of several of these key markers. A set of genes up-regulated in reset cells were also expressed in the human ICM and epiblast, and low or absent in various conventional and alternative primed PSC cultures (Fig. 4E; S4). These genes encode transcription factors, epigenetic regulators, metabolic components, and surface proteins, and provide several candidate markers of human naïve pluripotency. In addition we inspected recently published transcriptome data from Cynomolgus monkey embryos (Nakamura et al., 2016). Analysis of the most differentially expressed genes between reset and primed PSC separated the Cynomolgus samples into a pre-implantation and a post-implantation cluster, respectively (Fig. S5). Notably reset cells share features with the pre-implantation epiblast while primed PSC are more similar to pre-streak and gastrulating epiblast.

### Methylome status

Global DNA hypomethylation is a distinctive characteristic of mouse and human ICM cells (Guo et al., 2014; Lee et al., 2014; Smith et al., 2012) that is manifest in candidate human naïve PSC (Takashima et al., 2014; Theunissen et al., 2016). We performed whole genome bisulfite sequencing (BS-seq) on primed S6EOS and on reset S6EOS and H9 derived from independent experiments with or without addition of XAV. Methylation profiles were compared with previous datasets for primed PSC, human ICM cells (Guo et al., 2014), transgene reset PSC (H9-NK2; Takashima et al., 2014) and HNES1 cells (Guo et al., 2016). Primed PSC show uniform high levels of DNA methylation (85-95%), whereas reset cells display globally reduced CpG methylation, comparable to ICM and with a similar relatively broad distribution (Fig. 5A). Hypomethylation extended over all genomic elements (Fig. S6B) and was lower in cells that had been through more than 10 passages in t2iLGö. Loss of methylation from primed to reset conditions was not uniform across the whole genome, however. Highly methylated (80-100% mCpG) regions in primed cells showed divergent demethylation to between 15-65% (Fig. 5A,B & S6C). The majority of promoters were lowly methylated in both primed and reset S6EOS cells (Fig. 5C), including most CGI-containing promoters. Among methylated promoters in primed PSC, many showed decreased methylation in reset cells in line with the global trend. However, we also identified a number of CGI and non-CGI promoters that gained methylation upon resetting (highlighted in red; >40% CpG methylation difference between primed and averaged reset cells). Gene ontology (GO) analysis of the genes associated with this group of promoters indicated enrichment for terms related to differentiation, development and morphogenesis (Fig. S6D). Transgene reset and HNES1 cells also showed significantly higher promoter methylation levels at these loci than their primed counterparts (Fig. 5D), suggesting that selective promoter methylation is a feature of naïve-like cells in t2iLGö. In contrast we observed that many, although not all, imprinted DMRs are demethylated in reset conditions (Fig. 5E), in line with previous findings (Pastor et al., 2016).

**Figure 5.**
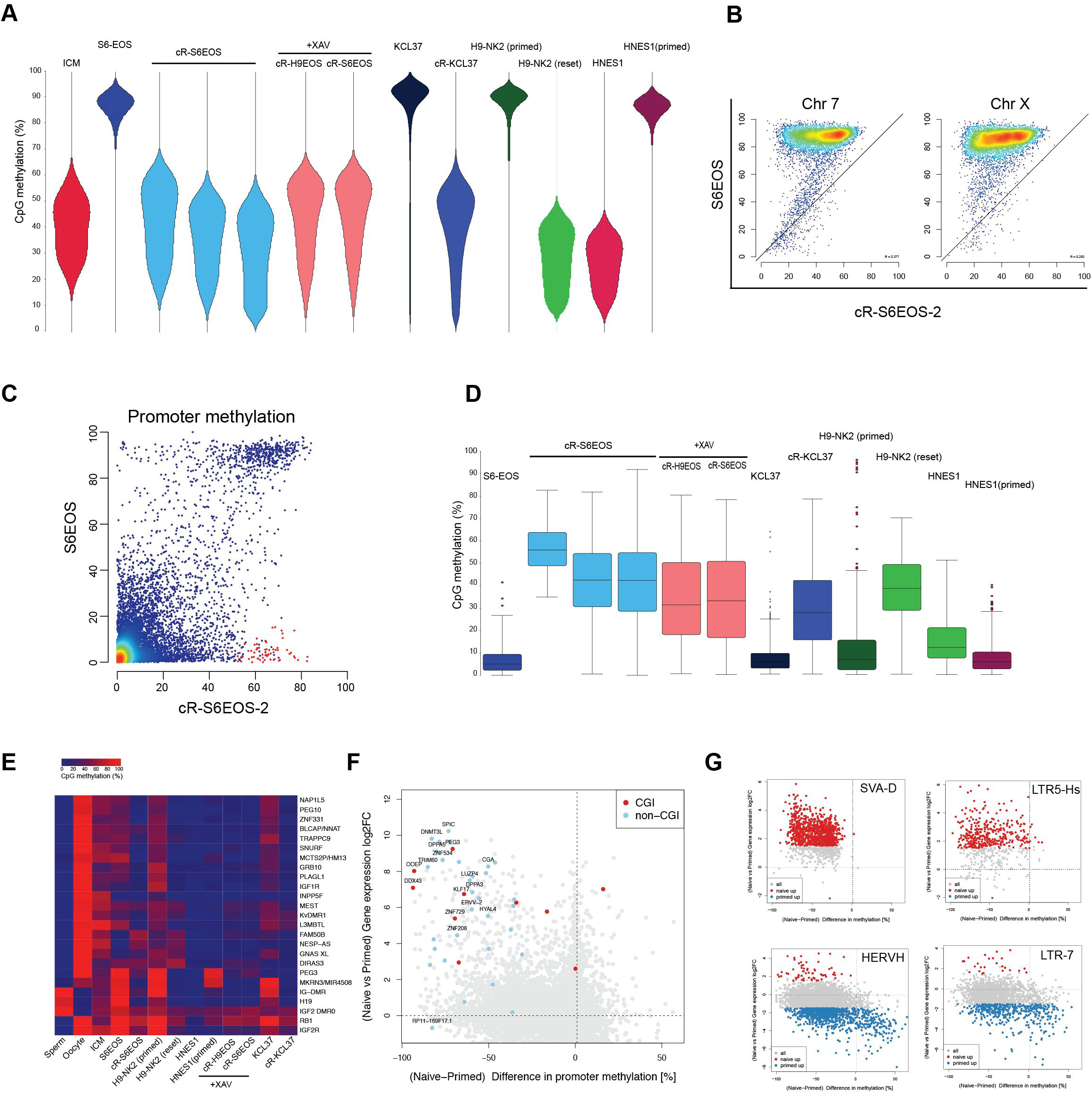
Methylome. A. Bean plots showing the global distribution of CpG methylation levels from pooled replicates of indicated samples compared with human ICM data (Guo 2014). Reset samples are from independent derivations without or with addition of XAV. Methylation was quantitated over 20 kb genomic tiles. Note KCL37 and HNES1 are male and H9 and Shef6 are female. B. Scatter plots of CpG methylation percentages over tiles spanning 20kb on Chromosome 7 and Chromosome X, comparing parental Shef6EOS in (KSR/FGF) with cR-S6EOS. C. Scatter plots of CpG methylation over promoters (-900 – 100), for parental and cRS6EOS cells. Promoters with > 40% gain in CpG methylation in reset cells are highlighted in red. D. CpG methylation levels of subset of promoters highlighted in panel C in indicated samples. E. Averaged CpG methylation of known DMRs of imprinted maternal and paternal genes. Sperm and oocyte data from (Okae et al., 2014); ICM from (Guo et al., 2014); H9 and H9-NK2 from (Takashima et al., 2014). F. Scatter plot showing the change in expression (log2FC) against the difference in promoter methylation for reset (averaged over cR-H9EOS and cR-S6EOS) versus parental Shef6EOS. G. Scatter plots for prominent differentially expressed transposon families showing the change in expression (log2FC) versus the difference in methylation for all loci.

The correlation between gene expression and promoter methylation (Fig. 5F, S6E) is very weak overall, as previously noted in mouse ES cells (Ficz et al., 2013; Habibi et al., 2013). Nonetheless, some genes that are highly up-regulated in reset cells and potentially functionally significant, such as *KLF17, DNMT3L* and *ZNF534*, show striking reductions in promoter methylation. Conversely, while TEs in general obeyed the genome-wide trend of hypomethylation in reset cells, substantial subsets of the HERVH and LTR7 TE families gained methylation and most of these showed reduced expression or were silenced (Fig. 5G). Finally, we noted demethylation of the piggyBac repeat sequences in cR-S6EOS cells (Fig. S6F), consistent with the proposition that the transgene is subject to epigenetic repression in primed cells that is relieved by resetting.

### Chromosomal stability

A major concern with manipulation of PSC culture conditions is the potential for selection of genetic variants (Amps et al., 2011). Indeed it has previously been noted that naïve-like cells cultured in the 5i/L/A formulation are prone to aneuploidy (Pastor et al., 2016; Sahakyan et al., 2017; Theunissen et al., 2014). We therefore carried out metaphase chromosome analyses by G-banding on a selection of cR cells (Fig. S2E). The results presented in Table 1 show retention of a diploid karyotype in most cases although in some cultures minor sub-populations of aneuploid cells are present. These data indicate that the epigenetic resetting process does not induce major chromosomal instability nor select for pre-existing variants, in line with previous observations that cultures in t2iLGö can maintain a diploid karyotype (Guo et al., 2016; Takashima et al., 2014). However, we noticed a variable incidence of tetraploid cells during expansion and one line showed a ubiquitous gain of chr19q13 after extended culture (40 passages). cR and HNES1 cells could also maintain a diploid karyotype over multiple passages in Geltrex or laminin, although abnormalities emerged in some cultures (Table 1). We also examined the transcriptome data by variant analysis for mutations in *TP53* which have been detected recurrently in primed PSC (Merkle et al., 2017). None of the loss-of-function *P53* mutations identified were found in cR cells.

### Differentiation

To assess the multi-lineage potential of chemically reset cells we first used embryoid body differentiation. After three days of floating culture in t2iL, aggregates were transferred to Geltrex-coated dishes and differentiated as outgrowths in serum. Alternatively reset cells were transferred into E8 for 6 days then aggregated in serum for 3 days before outgrowth. RT-qPCR on 8 day outgrowths showed up-regulation in both conditions of markers of early neuroectoderm, mesoderm and endoderm specification (Fig. S7A). Induction of these markers was lower for reset cells taken directly from t2iLGö than for cells conditioned in E8 (Fig. S7A), while down-regulation of pluripotency markers was similar. Immunostaining evidenced expression of protein markers of mesoderm and endoderm differentiation (Fig. S7B) and at lower frequency of neuron-specific beta-tubulin.

We then evaluated directed lineage commitment in adherent culture. Unsurprisingly, cR cells taken directly from t2iLGö did not respond directly to definitive endoderm or neuroectoderm induction protocols (Chambers et al., 2009; Loh et al., 2014) developed for primed PSC (Fig. S7C). After prior transfer into N2B27 for three days, a CXCR4/SOX17 positive, PDGFRα-negative, population, indicative of definitive endoderm, could be obtained (Fig. S7D) but neural marker induction in response to dual SMAD inhibition remained low. We therefore converted cR cells into a conventional primed PSC state by culture in E8 medium on Geltrex for several passages (Fig. S7E). We then applied the protocols for germ layer specification from primed cells to three different “re-primed” cultures. We observed robust expression of lineage markers for endoderm, lateral plate mesoderm and neuroectoderm by RT-qPCR (Fig. 6A). Immunostaining for SOX17 and FOXA2, and for SOX1 and PAX6, validated widespread generation of endoderm or neuroectoderm, respectively (Fig. 6B). Flow cytometric analysis quantified efficient induction of all three lineages (Fig. 6C, Fig. S7F). We examined further neuronal differentiation. After 29 days we detected expression of neuronal markers by RT-qPCR (Fig. 6D). Many cells with neurite-like processes were immunopositive for MAP2 and NeuN (Fig. 6E). By 40 days, markers of maturing neurons were apparent; vesicular glutamate transporter (vGlut2), post-synaptic protein 25 (SNAP25) and pre-synaptic protein bassoon (Fig. 6F).

**Figure 6.**
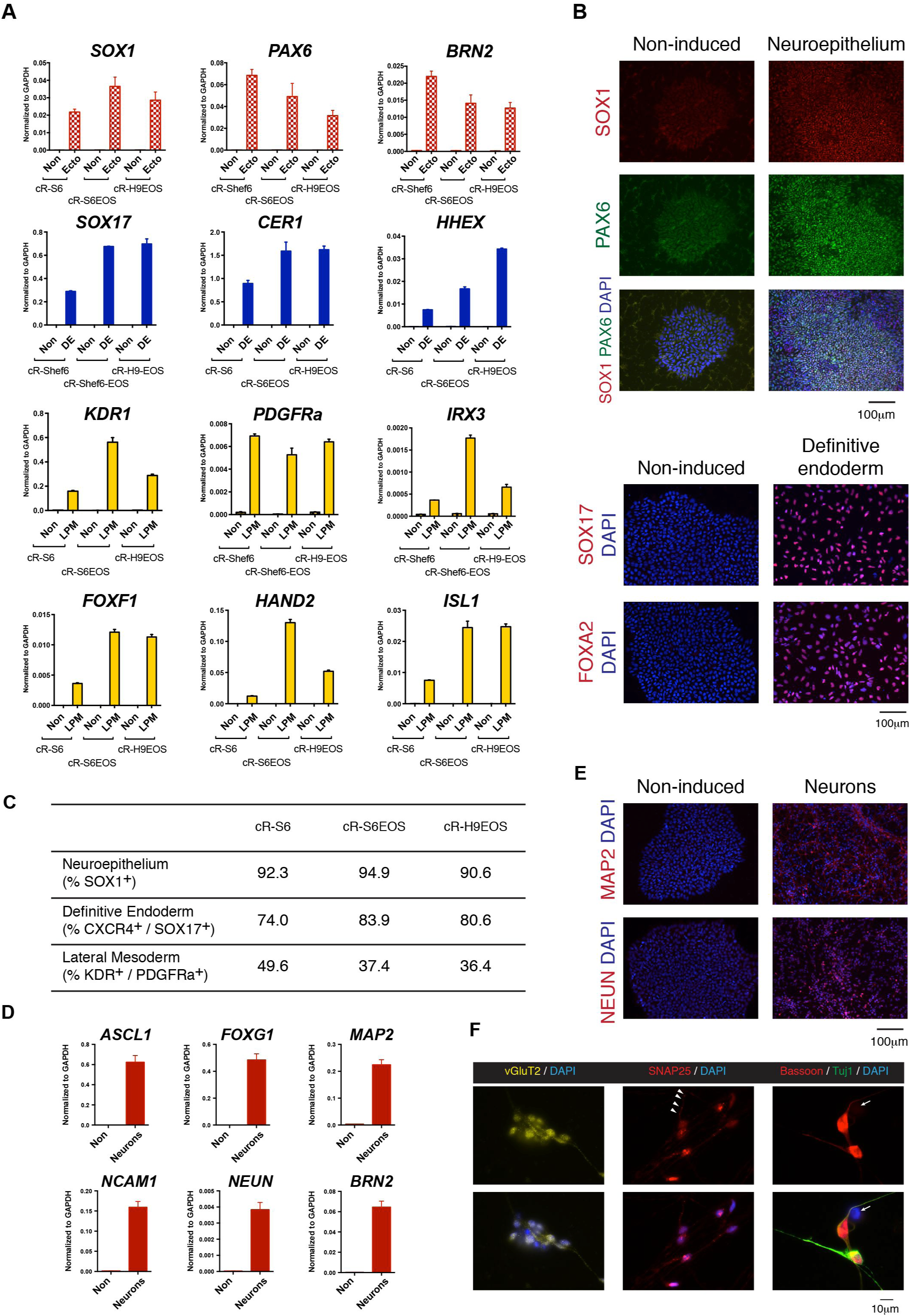
Differentiation. A. RT-qPCR analysis of lineage specification markers after induction of re-primed cR lines. Non indicates non-induced. B. Immunostaining for lineage specification markers. Scale bar 100 μM. C. Table summarizing flow cytometric quantification of neuroectodermal, mesodermal and endodermal lineage specification. D. RT-qPCR assays for pan-neuronal markers after 29 days differentiation from re-primed cR-S6EOS cells. E. Immunostaining for neuronal markers MAP2 and NEUN after 29 days. Scale bar 100 μM F. Immunostaining for neuronal maturation markers after 40 days. Arrowheads in middle panel highlight expected punctate clusters of SNAP25. Scale bar 20 μM

We also subjected cR-S6EOS cells to a protocol for inducing primordial germ cell-like cells (PGCLC). Cells were transferred from t2iLGö into TGFβ and FGF for 5 days, followed by exposure to germ cell inductive cytokines (Irie et al., 2015; von Meyenn et al., 2016). Cells co-expressing tissue non-specific alkaline phosphatase and EOS-GFP, suggestive of germ cell identity, were isolated by flow cytometry on day 9. Analysis of this double positive population by RT-qPCR showed up-regulated expression of a panel of PGC markers (Fig. S7G). These data indicate that germ cell specification may be induced from chemically reset cells, as recently also shown for reset cells generated by transgene expression (von Meyenn et al., 2016).

### X chromosome activity

Female naïve cells are expected to have two active X chromosomes in human as in mouse. Unlike in mouse, however, XIST is expressed from one or both active X chromosomes in human ICM cells (Okamoto et al., 2011; Petropoulos et al., 2016; Vallot et al., 2017) as well as from the inactive X in differentiated cells. Primed female hPSC usually feature an inactive X, although this has frequently lost XIST expression, a process referred to as erosion (Mekhoubad et al., 2012; Silva et al., 2008). X chromosomes in female cR-S6EOS cells show more marked loss of methylation than autosomes (Fig S6C), suggestive of reactivation (Takashima et al., 2014). We employed RNA-FISH to assess nascent transcription from X chromosomes at the single cell level. In parental S6EOS and H9EOS cells the presence of two X chromosomes was confirmed by RNA-FISH for *XACT* (Fig. S8A), which is transcribed from both active and eroded X chromosomes (Patel et al., 2017; Vallot et al., 2017). No *XIST* signal was evident in either cell line but we detected mono-allelic transcription of *HUWE1*, an X-linked gene typically subject to X chromosome inactivation (Patel et al., 2017)(Fig. 7A,B). In contrast, reset cells displayed bi-allelic transcription of *HUWE1* in the majority (90%) of diploid cells for both lines. Similar results were obtained for two other X-linked genes *ATRX* and *THOC2* (Fig.S8A,B). *XIST* was detected mono-allelically in a subset of reset cells (Fig 7A,B). This unusual feature is in line with recent reports that human naïve-like cells have two active X chromosomes, but predominantly express *XIST* from neither, or only one, allele (Sahakyan et al., 2017; Vallot et al., 2017).

**Figure 7.**
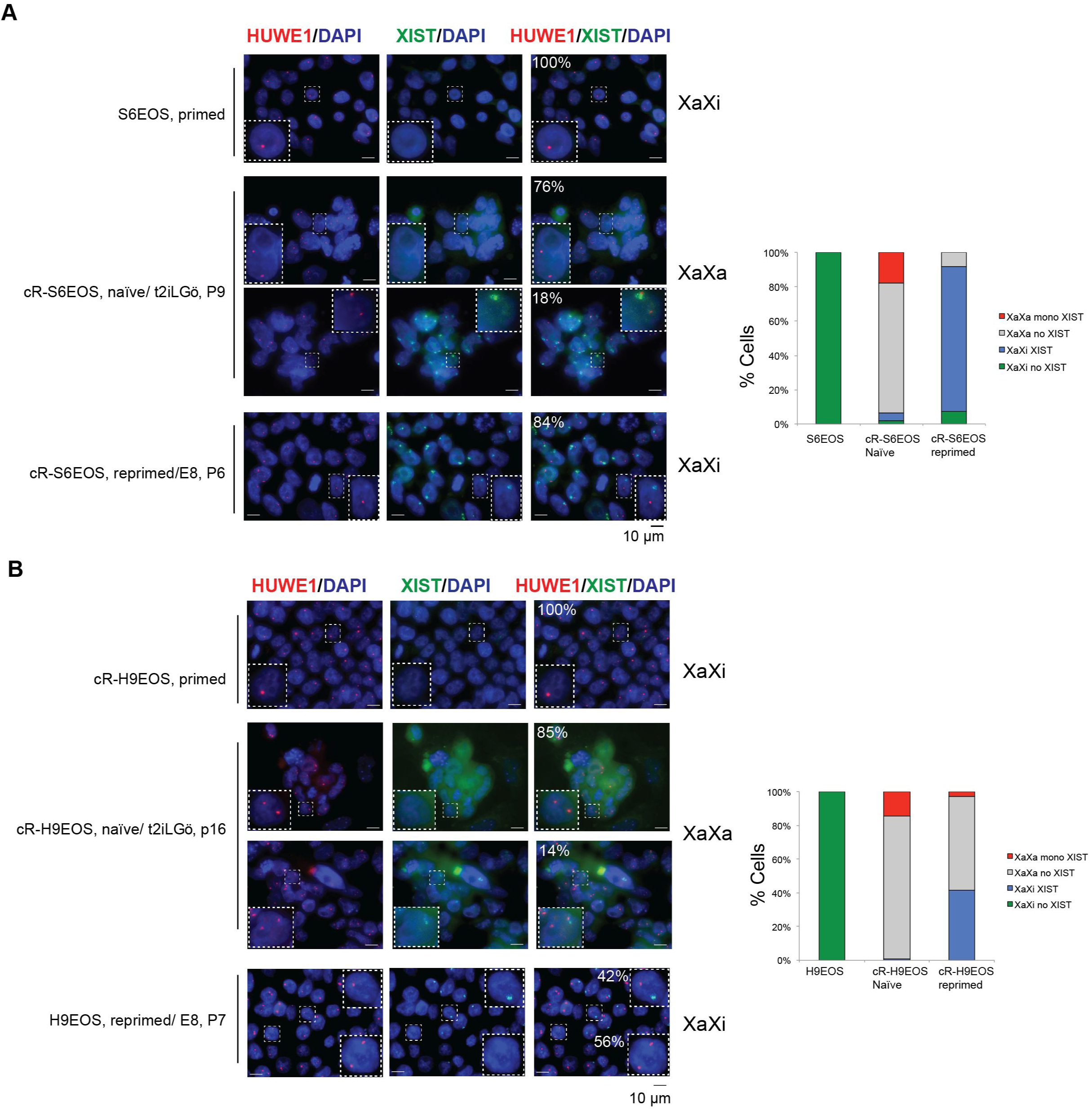
X chromosome status. RNA FISH for nascent X-linked RNA transcription in parental, reset and re-primed cells. A. S6EOS B. H9EOS Note that in re-primed cells displaying mono-allelic *HUWE1* and *XIST* expression, the two signals are on different chromosomes. Bar charts show quantification of X chromosome activation status based on *HUWE1* and *XIST* signals from samples of at least 100 cells.

We also examined X chromosome status after reset cells had been reverted to a primed-like PSC state by culture in E8 medium for 30 days as above. We found that *HUWE1* became transcribed monoallelically in around 90% of “re-primed” cR-S6EOS cells and that almost all of those cells expressed *XIST* from the other X (Fig. 7A,B). For cR-H9EOS, 40% of re-primed cells showed monoallelic expression of *HUWE1*, and those cells also up-regulated *XIST* from the other, inactive X. Similar patterns were observed when we co-stained the cells for *XIST* and another X-linked gene *THOC2* (Fig. S8A). These data are consistent with induction of X chromosome silencing by XIST during pluripotency progression.

## DISCUSSION

Availability of candidate naïve hPSC offers an experimental system for investigation of human pluripotency progression and a potentially valuable source material for biomedical applications. Our findings demonstrate that cell populations exhibiting a range of properties consistent with naive pluripotency can readily be generated from primed PSC by transient HDAC inhibition followed by culture in t2iLGö or tt2iLGö. WNT inhibition stabilises initial acquisition of the reset phenotype. Chemically reset cells are phenotypically stable and in many cases cytogenetically normal. They can be propagated robustly without feeders and readily be re-primed to undergo multi-lineage differentiation in vitro. We provide detailed protocols along with global transcriptome, transposcriptome, and methylome datasets as resources for the community.

The mechanism by which HDAC inhibition promotes resetting is unresolved but seems likely to involve generation of a more open chromatin environment that relieves silencing of naïve pluripotency factors. The reset phenotype is initially rather precarious but can be stabilised by inhibitors of tankyrase or porcupine that suppress the canonical WNT pathway. cR cells differ dramatically in global expression profile from primed PSC and resemble previously described human naïve-like cells generated by inducible or transient transgene expression (Takashima et al., 2014), or by adaptation to culture in 5i/L/A/(F) (Theunissen et al., 2014). In particular, transcriptome analysis shows that cR cells share a high degree of genomewide and marker-specific correspondence with HNES cell lines derived directly from dissociated human ICM (Guo et al., 2016). Reset cells express transcription regulators and other genes that are found in human pre-implantation epiblast but are low or absent in primed PSC. These include the characterised naïve pluripotency factors KLF4 and TFCP2L1 along with potential new regulators and markers.

Reset and HNES cells express SVA, LTR5, HERVK and SST1 transposable elements. These are among the most recent entrants to the human genome and are transcribed in pre-implantation embryos (Grow et al., 2015; Theunissen et al., 2016). In contrast HERVH families and their flanking LTR7 repeats are mostly down-regulated in reset cells and exhibit increased methylation. These findings confirm and extend the recent report that specific TE expression discriminates between primed and naïve-like human PSC (Theunissen et al., 2016). HERVH and LTR7 are reported to generate alternative and chimaeric transcripts in primed PSC where they display heterogeneous expression (Wang et al., 2014). Therefore, silencing in naïve cells and de-repression upon progression to primed pluripotency may have functional significance. Notably ZNF534, the postulated negative regulator of HERVH (Theunissen et al., 2016), is highly up-regulated in reset cells, while increased expression of DNMT3L in human naïve-like cells, a feature not apparent in mouse ESC, may facilitate de novo methylation at specific TE loci.

During resetting, DNA methylation is globally reduced to a level similar to that reported for human ICM (Guo et al., 2014). This is regarded as a key process for erasure of epigenetic memory in the naïve phase of pluripotency (Lee et al., 2014). Reduced methylation extends to all classes of genomic element but is non-uniform. At promoters, both loss and gain of methylation are detected. As in other cell types, there is poor overall correlation with gene expression but it is noteworthy that extensively demethylated promoters in reset cells include several associated with highly up-regulated genes that are likely to be functional in naïve cells, including *KLF17,* as well as numerous primate and hominid-specific transposable elements. Demethylation also extends to imprinted loci, however, as noted previously for other human naïve-like stem cells (Pastor et al., 2016; Theunissen et al., 2016). Loss of imprints is observed in conventional hPSC (Nazor et al., 2012) and in mouse ESC (Dean et al., 1998; Greenberg and Bourc’his, 2015; Walter et al., 2016), but not typically to the extent detected for human naïve-like cells. Whether failure to sustain imprints is an intrinsic feature of human naïve pluripotency during extended propagation or may be rectified by modification of the culture environment remains to be determined.

Efficient multi-lineage differentiation may be initiated from reset cells either via embryoid body formation, or by “re-priming” in adherent culture. It is noteworthy, however, that human cells in the t2iLGö naïve condition are not immediately responsive to lineage induction. Ground state mouse ES cells also appear not to respond directly to lineage cues but to require prior transition through a formative stage (Kalkan et al., 2017; Mulas et al., 2016; Semrau et al., 2016). This capacitation period may be more protracted in primates, given the longer window between implantation and gastrulation (Nakamura et al., 2016; Smith, 2017).

A hallmark of the transient phase of naïve pluripotency in both rodent and human ICM cells is the presence of two active X chromosomes in females (Okamoto et al., 2011; Petropoulos et al., 2016; Sahakyan et al., 2017; Vallot et al., 2017). In female chemically reset cells, the gain of biallelic expression of X-linked genes indicates reactivation of the silent X chromosome. Moreover, expression of *XIST* from an active X chromosome in a subset of reset cells resembles the pattern of the human pre-implantation embryo. Upon re-priming monoallelic expression of X-linked genes is restored in many cells. Significantly, although no XIST was observed in the original primed cells, an XIST signal is detected in re-primed cells on a silenced X chromosome. Resetting and subsequent differentiation thus offer a system to characterise X chromosome regulation in human, which appears to diverge substantially from the mouse paradigm (Okamoto et al., 2011).

In summary, this study provides the requisite technical protocols and resources to facilitate routine generation and study of candidate human naïve PSC. Moreover, feeder-free culture simplifies the propagation of reset cells. Nonetheless, further refinements are desirable to enhance the quality and robustness of human naïve pluripotent stem cells, including preserving imprints and maximising long-term karyotype stability. Optimising the capacitation process prior to differentiation by recapitulating the progression of pluripotency in the primate embryo is an important future goal and opportunity.

### Materials and methods

#### Conventional hPSC culture

Primed hPSC were routinely maintained on irradiated mouse embryo fibroblast feeder cells (MEFs) in KSR/FGF medium: DMEM/F-12 (Sigma-Aldrich, D6421) supplemented with 10 ng/ml FGF2 (prepared in-house), 20% KnockOut Serum Replacement (KSR) (Thermo Fisher Scientific), 100 mM 2-mercaptoethanol (2ME) (Sigma-Aldrich, M7522), 1XMEM nonessential amino acids (NEAA) (Thermo Fisher Scientific, 11140050) and 2 mM Lglutamine (Thermo Fisher Scientific, 25030024). Cells were passaged as clusters by detachment with dispase (Sigma-Aldrich, 11097113001). To establish *PB-Oct4EOS* stable transfectants, 1 μg/ml of puromycin was applied for two passages (10 days) to transfected cells on Matrigel (Roche). Some PSC lines were propagated without feeders on Geltrex (growth factor-reduced, Thermo Fisher, A1413302) in E8 medium (made in-house according to (Chen et al., 2011).

#### Naïve cell culture

Chemically reset and embryo derived (HNES) naïve stem cells were propagated in N2B27 (see Supplemental Protocol) supplemented with t2iLGö (1μM CHIR99021, 1μM PDO325901, 10ng/ml human LIF and 2μM Gö6983) with or without Rock inhibitor (Y-27632) on irradiated MEF feeders. Where indicated as tt2iLGö, CH was used at 0.3μM. For feeder-free culture, Geltrex or laminin (Merck, CC095) were added to the medium at time of plating. Cells were cultured in 5% O2, 7% CO2 in a humidified incubator at 37°C and passaged by dissociation with Accutase (Thermo Fisher Scientific, A1110501) or TrypLE (Thermo Fisher Scientific, 12605028) every 3-5 days. Cells were cryopreserved in CryoStem (Biological Industries, K1-0640). All cell lines were tested free of mycoplasma contamination in lab by PCR. No other contamination test has been performed.

#### Reverse transcription and real-time PCR

Total RNA was extracted using RNAeasy (Quiagen) and cDNA synthesized with SuperScript^®^ III Reverse Transcriptase and Oligo-dT adapter primers. Taqman assays and UPL probes are listed in Table S4A and Table S4B. Embryoid bodies were lysed in TRIzol and total RNA was isolated with PureLink RNA Mini Kit (Thermo Fisher Scientific, 12183025) with On-Column PureLink DNAase (Thermo Fisher Scientific, 12185010). For analyses of adherent differentiation, total RNA was extracted with Reliaprep RNA Miniprep kit and RTqPCR performed using Oligo-dT primer, Goscript Reverse Transcription system and GoTaq qPCR Master Mix (all from Promega).

#### Immunostaining

Cells were fixed with 4% buffered formaldehyde for 15min at room temperature, permeabilised with 0.5% Triton X-100 in PBS for 10min and blocked with 3% BSA and 0.1% Tween-20 in PBS for 30min at room temperature. Incubation with primary antibodies (Table S4C) diluted in PBS with 0.1% Triton X-100 and 3% donkey serum was overnight at +4°C and secondary antibodies were added for 1 hour at room temperature. Slides were mounted with Prolong Diamond Antifade Mountant (Life Technologies)

#### Chromosome analysis

G-banded karyotype analysis was performed following standard cytogenetics protocols, typically scoring 30 metaphases.

#### Transcriptome sequencing

Total RNA was extracted using TRIzol/chloroform method (Invitrogen) and RNA integrity assessed by Qubit measurement and RNA nanochip Bioanalyzer. Ribosomal RNA was depleted from 1 pg of total RNA using Ribozero (Illumina Kit). Sequencing libraries were prepared using the NEXTflex Rapid Directional RNA-Seq Kit. Sequencing was performed on the Illumina HiSeq4000 in either single end 50bp or paired end 125bp formats.

#### RNA-seq data analysis

External datasets used for comparative analyses were obtained from the European Nucleotide Archive (ENA) under accessions ERP006823 (Takashima et al., 2014), SRP059279 (Ji et al., 2016), SRP045911 (Sperber et al., 2015), SRP045294 (Irie et al., 2015), and ERP007180 (Wellcome Trust Sanger Institute). To minimise technical variability reads of disparate lengths and sequencing modes were truncated to 50bp single-end format. Alignments to human genome build hg38/GRCh38 were performed with STAR (Dobin et al., 2013). Transcript quantification was performed with htseq-count, part of the HTSeq package (Anders et al., 2014), using gene annotation from Ensembl release 86 (Aken et al., 2016). Libraries were corrected for total read count using the size factors computed by the Bioconductor package DESeq2 (Love et al., 2014). Principal components were computed by singular value decomposition with the prcomp function in the R stats package from variance-stabilized count data. Differential expression was computed with DESeq2 and genes ranked by log_2_ fold change. t-Distributed stochastic neighbor embedding (t-SNE) (van der Maaten and Hinton, 2008) was performed using the Barnes-Hut algorithm (Van Der Maaten, 2014) implemented in the Bioconductor package Rtsne with perplexity 12 for 1600 iterations. For display of expression values, single-end count data were normalised for gene length to yield RPKMs and scaled relative to the mean expression of each gene across all samples. Heatmaps include genes for which a difference in expression was observed (i.e., scaled expression > 1 or < -1 in at least one sample). For functional testing, enrichment for Gene Ontology (GO) terms was found using the GOStats package (Falcon & Gentleman, 2007) based on the 1000 most up- and downregulated genes distinguishing naïve and primed cells, and most significant genes contributing to principal component 1 (Fig 3A). RNA-seq libraries were screened for mutations in the P53 locus by processing alignments with Picard tools (http://broadinstitute.github.io/picard) and the Genome Analysis Toolkit (GATK)(DePristo et al., 2011; McKenna et al., 2010) to filter duplicate reads, perform base quality score recalibration, identify indels for realignment, and call variants against dbSNP Build 150 (Sherry et al., 2001).

#### Bisulfite Sequencing, Mapping and Analysis

Post-bisulfite adaptor tagging (PBAT) libraries for whole-genome DNA methylation analysis were prepared from purified genomic DNA (Miura et al., 2012; Smallwood et al., 2014; von Meyenn et al., 2016). Paired-end sequencing was carried out on HiSeq2000 or NextSeq500 instruments (Illumina). Raw sequence reads were trimmed to remove poor quality reads and adapter contamination using Trim Galore (v0.4.1). The remaining sequences were mapped using Bismark (v0.14.4) (Krueger and Andrews, 2011) to the human reference genome GRCh37 in paired-end mode as described (von Meyenn et al., 2016). CpG methylation calls were analyzed using SeqMonk software and custom R scripts. Global CpG methylation levels of pooled replicates were illustrated using bean plots. The genome was divided into consecutive 20 kb tiles and percentage methylation was calculated using the bisulfite feature methylation pipeline in SeqMonk. Pseudocolor scatter plots of methylation levels over 20 kb tiles were generated using R.

Specific genome features were defined using the Ensembl gene sets annotations: Gene bodies (probes overlapping genes), Promoters (probes overlapping 900bp upstream to 100bp downstream of genes), CGI promoters (promoters containing a CGI), non-CGI promoters (all other promoters), Intergenic (probes not overlapping with gene bodies), non-promoter CGI (CGI not overlapping with promoters). Annotations of human germline imprint control regions were obtained from (Court et al., 2014). Pseudocolor heatmaps representing average methylation levels were generated using the R “heatmap.2” function without further clustering, scaling or normalization. Correlation between promoter methylation and gene expression was computed from average CpG methylation across promoters or transposable elements and correlating these values with the respective gene expression values.

#### Fluorescent in situ hybridization (FISH)

Nascent transcription foci of X-linked genes and the lncRNAs XIST and XACT were visualized at single-cell resolution by RNA FISH as described (Sahakyan et al., 2017). Fluorescently labeled probes were generated from BACs RP11-13M9 (*XIST)*, RP11-35D3 (*XACT*), RP11-121P4 (THOC2), RP11-1145J4 (*ATRX*), and RP11-975N19 (*HUWE1*). Coverslips were imaged using an Imager M1 microscope (Zeiss) and Axio Vision software. ImageJ was used for collapsing Z-stacks, merging different channels, and adjusting brightness and contrast to remove background. A minimum of 100 nuclei were scored for each sample. Cells that appeared to have more than two X chromosomes were excluded.

#### Transposable elements

RepeatMasker annotations for the human reference genome were obtained from the UCSC Table Browser. To calculate repeat expression, adapter-trimmed RNA-seq reads were mapped to the reference genome by using bowtie (<u>http://bowtiebio.sourceforge.net</u>; version: 1.1.0) with parameters ‘-M1 –v2 –best –strata’; i.e. two mismatches were allowed, and one alignment location was randomly selected for reads that multiply align to the reference genome. Read counts for repeat regions and ENSEMBL transcripts were calculated by featureCounts, normalised by the total number of RNA-seq reads that mapped to protein-coding gene regions. Differential expression of repeat copies across samples was evaluated by the R Bioconductor DESeq package (Anders and Huber, 2010).

#### Embryoid body differentiation

Embryoid body formation and outgrowth were performed in DMEM/F12 supplemented with 15% FCS, 2mM L-glutamine. 1mM sodium pyruvate, 1x non-essential amino acids and 0.1 mM 2-mercaptoethanol as described (Guo et al., 2016). Alternatively reset cells were aggregated in t2iLIF medium with ROCKi in PrimeSurface 96V cell plates (Sumitomo Bakelite MS-9096V) then plated after three days on Geltrex (Thermo Fisher Scientific, 12063569) for outgrowth in serum-containing medium. Outgrowths were fixed with 4% PFA for 10 minutes at room temperature for immunostaining.

#### Adherent differentiation

Except where specified, reset cells were “re-primed” before initiating differentiation. Cells were plated on Geltrex in t2iLGö and after 48 hours the medium was changed to E8. Cultures were maintained in E8, passaging at confluency. Lineage-specific differentiation was initiated between 25-44 days.

Definitive endoderm was induced according to (Loh et al., 2014). Cells were cultured in CDM2 medium supplemented with 100 ng/ml Activin A (produced in-house), 100 nM PI-103 (Bio-techne, 2930), 3 µM CHIR99021, 10 ng/ml FGF2, 3 ng/ml BMP4 (Peprotech) for one day. For the next 2 days the following supplements were applied: 100ng/ml Activin A, 100nM PI-103, 20ng/ml FGF2, 250nM LDN193189.

For lateral mesoderm induction (Loh et al., 2016) cells were treated with CDM2 supplemented with 30ng/ml Activin A, 40ng/ml BMP4 (Miltenyi Biotech, 130-098-788), 6μM CHIR99021, 20ng/ml FGF2, 100nM PI-103 for 1 day, then with 1μM A8301, 30ng/ml BMP4 and 10μM XAV939 (Sigma-Aldrich).

For neural differentiation via dual Smad inhibition (Chambers et al., 2009), cells were treated with N2B27 medium supplemented with 500 nM LDN193189 (Axon, 1509) and 1 μM A 83- 01 (Bio-techne, 2939) for 8-10 days), then passaged to plates coated with poly-ornithine and laminin and further cultured in N2B27 without supplements.

#### Flow cytometry

Flow analysis was carried out on a Fortessa instrument. Cell sorting was performed using a MoFlo high speed instrument.

## Acknowledgements

Rosalind Drummond provided excellent technical support. We thank Nicholas Bredenkamp for sharing data. We are grateful to Peter Andrews for advice and support on karyotyping and to Valeria Orlova and Balazs Varga for advice on differentiation protocols. Andy Riddell and Peter Humphreys supported flow cytometry and imaging studies. Maike Paramor prepared RNA-seq libraries. Sequencing was conducted at the CRUK Cambridge Institute Genomic Core.

## Author Contributions

GG and ASm designed the study. GG, MR, JC and SM performed experiments and prepared samples; DB analysed metaphase chomosomes; ASa performed FISH studies, overseen by KP; FvM carried out methylome analyses, overseen by WR; SD and PB analysed transcriptome data. ASm and GG wrote the paper with input from other authors.

## Data availability

RNA-seq data are deposited in ArrayExpress under accession number E-MTAB-5674, WGBS data in GEO under accession number GSE90168.

## References

Amps, K. Andrews, P. W. Anyfantis, G. Armstrong, L. Avery, S. Baharvand, H. Baker, J. Baker, D. Munoz, M. B. Beil, S. et al. (2011). Screening ethnically diverse human embryonic stem cells identifies a chromosome 20 minimal amplicon conferring growth advantage. Nat Biotechnol 29, 1132–44.

Anders, S. and Huber, W. (2010). Differential expression analysis for sequence count data. Genome Biol 11, R106.

Blakeley, P., Fogarty, N. M. E., del Valle, I., Wamaitha, S. E., Hu, T. X., Elder, K., Snell, P., Christie, L., Robson, P. and Niakan, K. K. (2015). Defining the three cell lineages of the human blastocyst by single-cell RNA-seq. Development.

Boroviak, T., Loos, R., Bertone, P., Smith, A. and Nichols, J. (2014). The ability of innercell-mass cells to self-renew as embryonic stem cells is acquired following epiblast specification. Nat Cell Biol 16, 516–28.

Boroviak, T., Loos, R., Lombard, P., Okahara, J., Behr, R., Sasaki, E., Nichols, J., Smith, A. and Bertone, P. (2015). Lineage-Specific Profiling Delineates the Emergence and Progression of Naive Pluripotency in Mammalian Embryogenesis. Developmental Cell 35, 366–382.

Brons, I. G., Smithers, L. E., Trotter, M. W., Rugg-Gunn, P., Sun, B., Chuva de Sousa Lopes, S. M., Howlett, S. K., Clarkson, A., Ahrlund-Richter, L., Pedersen, R. A. et al. (2007). Derivation of pluripotent epiblast stem cells from mammalian embryos. Nature 448, 191–5.

Butcher, L. M., Ito, M., Brimpari, M., Morris, T. J., Soares, F. A., Ahrlund-Richter, L., Carey, N., Vallier, L., Ferguson-Smith, A. C. and Beck, S. (2016). Non-CG DNA methylation is a biomarker for assessing endodermal differentiation capacity in pluripotent stem cells. Nat Commun 7, 10458.

Chambers, S. M., Fasano, C. A., Papapetrou, E. P., Tomishima, M., Sadelain, M. and Studer, L. (2009). Highly efficient neural conversion of human ES and iPS cells by dual inhibition of SMAD signaling. Nat Biotechnol 27, 275–80.

Chen, B., Dodge, M. E., Tang, W., Lu, J., Ma, Z., Fan, C. W., Wei, S., Hao, W., Kilgore, J., Williams, N. S. et al. (2009). Small molecule-mediated disruption of Wnt-dependent signaling in tissue regeneration and cancer. Nat Chem Biol 5, 100–7.

Chen, G., Gulbranson, D. R., Hou, Z., Bolin, J. M., Ruotti, V., Probasco, M. D., Smuga-Otto, K., Howden, S. E., Diol, N. R., Propson, N. E. et al. (2011). Chemically defined conditions for human iPSC derivation and culture. Nat Methods 8, 424–9.

Court, F., Tayama, C., Romanelli, V., Martin-Trujillo, A., Iglesias-Platas, I., Okamura, K., Sugahara, N., Simon, C., Moore, H., Harness, J. V. et al. (2014). Genome-wide parent-of-origin DNA methylation analysis reveals the intricacies of human imprinting and suggests a germline methylation-independent mechanism of establishment. Genome Res 24, 554–69.

D’Aniello, C., Fico, A., Casalino, L., Guardiola, O., Di Napoli, G., Cermola, F., De Cesare, D., Tate, R., Cobellis, G., Patriarca, E. J. et al. (2015). A novel autoregulatory loop between the Gcn2-Atf4 pathway and L-Proline metabolism controls stem cell identity. Cell Death Differ 22, 1094–1105.

Davidson, K. C., Mason, E. A. and Pera, M. F. (2015). The pluripotent state in mouse and human. Development 142, 3090–3099.

De Los Angeles, A., Loh, Y. H., Tesar, P. J. and Daley, G. Q. (2012). Accessing naive human pluripotency. Curr Opin Genet Dev 22, 272–82.

Dean, W., Bowden, L., Aitchison, A., Klose, J., Moore, T., Meneses, J. J., Reik, W. and Feil, R. (1998). Altered imprinted gene methylation and expression in completely ES cell-derived mouse fetuses; association with aberrant phenotypes. Development 125, 2273–2282.

DePristo, M. A., Banks, E., Poplin, R., Garimella, K. V., Maguire, J. R., Hartl, C., Philippakis, A. A., del Angel, G., Rivas, M. A., Hanna, M. et al. (2011). A framework for variation discovery and genotyping using next-generation DNA sequencing data. Nat Genet 43, 491–8.

Ficz, G., Hore, T. A., Santos, F., Lee, H. J., Dean, W., Arand, J., Krueger, F., Oxley, D., Paul, Y. L., Walter, J. et al. (2013). FGF signaling inhibition in ESCs drives rapid genomewide demethylation to the epigenetic ground state of pluripotency. Cell Stem Cell 13, 351–9.

Friedli, M. and Trono, D. (2015). The Developmental Control of Transposable Elements and the Evolution of Higher Species. Annu Rev Cell Dev Biol 31, 429–451.

Gafni, O., Weinberger, L., Mansour, A. A., Manor, Y. S., Chomsky, E., Ben-Yosef, D., Kalma, Y., Viukov, S., Maza, I., Zviran, A. et al. (2013). Derivation of novel human ground state naive pluripotent stem cells. Nature 504, 282–6.

Greenberg, M. V. and Bourc’his, D. (2015). Cultural relativism: maintenance of genomic imprints in pluripotent stem cell culture systems. Curr Opin Genet Dev 31, 42–9.

Grow, E. J., Flynn, R. A., Chavez, S. L., Bayless, N. L., Wossidlo, M., Wesche, D. J., Martin, L., Ware, C. B., Blish, C. A., Chang, H. Y. et al. (2015). Intrinsic retroviral reactivation in human preimplantation embryos and pluripotent cells. Nature.

Guo, G., von Meyenn, F., Santos, F., Chen, Y., Reik, W., Bertone, P., Smith, A. and Nichols, J. (2016). Naive Pluripotent Stem Cells Derived Directly from Isolated Cells of the Human Inner Cell Mass. Stem Cell Reports 6, 437–46.

Guo, H., Zhu, P., Yan, L., Li, R., Hu, B., Lian, Y., Yan, J., Ren, X., Lin, S., Li, J. et al. (2014). The DNA methylation landscape of human early embryos. Nature 511, 606–610.

Habibi, E., Brinkman, A. B., Arand, J., Kroeze, L. I., Kerstens, H. H., Matarese, F., Lepikhov, K., Gut, M., Brun-Heath, I., Hubner, N. C. et al. (2013). Whole-genome bisulfite sequencing of two distinct interconvertible DNA methylomes of mouse embryonic stem cells. Cell Stem Cell 13, 360–9.

Hackett, J. A. and Surani, M. A. (2014). Regulatory principles of pluripotency: from the ground state up. Cell Stem Cell 15, 416–30.

Hotta, A., Cheung, A. Y., Farra, N., Vijayaragavan, K., Seguin, C. A., Draper, J. S., Pasceri, P., Maksakova, I. A., Mager, D. L., Rossant, J. et al. (2009). Isolation of human iPS cells using EOS lentiviral vectors to select for pluripotency. Nat Methods 6, 370–6.

Huang, K., Maruyama, T. and Fan, G. (2014). The naive state of human pluripotent stem cells: a synthesis of stem cell and preimplantation embryo transcriptome analyses. Cell Stem Cell 15, 410–5.

Huang, S.-M. A., Mishina, Y. M., Liu, S., Cheung, A., Stegmeier, F., Michaud, G. A., Charlat, O., Wiellette, E., Zhang, Y., Wiessner, S. et al. (2009). Tankyrase inhibition stabilizes axin and antagonizes Wnt signalling. Nature 461, 614–620.

Huangfu, D., Maehr, R., Guo, W., Eijkelenboom, A., Snitow, M., Chen, A. E. and Melton, D. A. (2008). Induction of pluripotent stem cells by defined factors is greatly improved by small-molecule compounds. Nat Biotechnol 26, 795–7.

Irie, N., Weinberger, L., Tang, W. W., Kobayashi, T., Viukov, S., Manor, Y. S., Dietmann, S., Hanna, J. H. and Surani, M. A. (2015). SOX17 is a critical specifier of human primordial germ cell fate. Cell 160, 253–68.

Ji, X., Dadon, D. B., Powell, B. E., Fan, Z. P., Borges-Rivera, D., Shachar, S., Weintraub, A. S., Hnisz, D., Pegoraro, G., Lee, T. I. et al. (2016). 3D Chromosome Regulatory Landscape of Human Pluripotent Cells. Cell Stem Cell 18, 262–75.

Kalkan, T., Olova, N., Roode, M., Mulas, C., Lee, H. J., Nett, I., Marks, H., Walker, R., Stunnenberg, H. G., Lilley, K. S. et al. (2017). Tracking the embryonic stem cell transition from ground state pluripotency. Development.

Kalkan, T. and Smith, A. (2014). Mapping the route from naive pluripotency to lineage specification. Phil Trans R Soc B 369.

Kim, H., Wu, J., Ye, S., Tai, C. I., Zhou, X., Yan, H., Li, P., Pera, M. and Ying, Q. L. (2013). Modulation of beta-catenin function maintains mouse epiblast stem cell and human embryonic stem cell self-renewal. Nat Commun 4, 2403.

Krueger, F. and Andrews, S. R. (2011). Bismark: a flexible aligner and methylation caller for Bisulfite-Seq applications. Bioinformatics 27, 1571–2.

Lee, H. J., Hore, T. A. and Reik, W. (2014). Reprogramming the methylome: erasing memory and creating diversity. Cell Stem Cell 14, 710–9.

Loh, K. M., Ang, L. T., Zhang, J., Kumar, V., Ang, J., Auyeong, J. Q., Lee, K. L., Choo, S. H., Lim, C. Y., Nichane, M. et al. (2014). Efficient endoderm induction from human pluripotent stem cells by logically directing signals controlling lineage bifurcations. Cell Stem Cell 14, 237–52.

Loh, K. M., Chen, A., Koh, P. W., Deng, T. Z., Sinha, R., Tsai, J. M., Barkal, A. A., Shen, K. Y., Jain, R., Morganti, R. M. et al. (2016). Mapping the Pairwise Choices Leading from Pluripotency to Human Bone, Heart, and Other Mesoderm Cell Types. Cell 166, 451–67.

Martello, G. and Smith, A. (2014). The nature of embryonic stem cells. Annu Rev Cell Dev Biol 30, 647–75.

McKenna, A., Hanna, M., Banks, E., Sivachenko, A., Cibulskis, K., Kernytsky, A., Garimella, K., Altshuler, D., Gabriel, S., Daly, M. et al. (2010). The Genome Analysis Toolkit: a MapReduce framework for analyzing next-generation DNA sequencing data. Genome Res 20, 1297–303.

Mekhoubad, S., Bock, C., de Boer, A. S., Kiskinis, E., Meissner, A. and Eggan, K. (2012). Erosion of dosage compensation impacts human iPSC disease modeling. Cell Stem Cell 10, 595–609.

Merkle, F. T., Ghosh, S., Kamitaki, N., Mitchell, J., Avior, Y., Mello, C., Kashin, S., Mekhoubad, S., Ilic, D., Charlton, M. et al. (2017). Human pluripotent stem cells recurrently acquire and expand dominant negative P53 mutations. Nature In press.

Miura, F., Enomoto, Y., Dairiki, R. and Ito, T. (2012). Amplification-free whole-genome bisulfite sequencing by post-bisulfite adaptor tagging. Nucleic Acids Res 40, e136.

Mulas, C., Kalkan, T. and Smith, A. (2016). Nodal secures pluripotency upon embryonic stem cell progression from the ground state. bioRxiv.

Nakamura, T., Okamoto, I., Sasaki, K., Yabuta, Y., Iwatani, C., Tsuchiya, H., Seita, Y., Nakamura, S., Yamamoto, T. and Saitou, M. (2016). A developmental coordinate of pluripotency among mice, monkeys and humans. Nature 537, 57–62.

Nazor, Kristopher L., Altun, G., Lynch, C., Tran, H., Harness, Julie V., Slavin, I., Garitaonandia, I., Müller, F.-J., Wang, Y.-C., Boscolo, Francesca S. et al. (2012). Recurrent Variations in DNA Methylation in Human Pluripotent Stem Cells and Their Differentiated Derivatives. Cell Stem Cell 10, 620–634.

Nichols, J. and Smith, A. (2009). Naive and primed pluripotent states. Cell Stem Cell 4, 487–92.

Nichols, J. and Smith, A. (2012). Pluripotency in the embryo and in culture. CSH Perspect Biol 4, a008128.

Nishizawa, M., Chonabayashi, K., Nomura, M., Tanaka, A., Nakamura, M., Inagaki, A., Nishikawa, M., Takei, I., Oishi, A., Tanabe, K. et al. (2016). Epigenetic Variation between Human Induced Pluripotent Stem Cell Lines Is an Indicator of Differentiation Capacity. Cell Stem Cell.

Ogura, A., Inoue, K. and Wakayama, T. (2013). Recent advancements in cloning by somatic cell nuclear transfer. Philos Trans R Soc Lond B Biol Sci 368, 20110329.

Okamoto, I., Patrat, C., Thepot, D., Peynot, N., Fauque, P., Daniel, N., Diabangouaya, P., Wolf, J. P., Renard, J. P., Duranthon, V. et al. (2011). Eutherian mammals use diverse strategies to initiate X-chromosome inactivation during development. Nature 472, 370–4.

Pastor, W. A., Chen, D., Liu, W., Kim, R., Sahakyan, A., Lukianchikov, A., Plath, K., Jacobsen, S. E. and Clark, A. T. (2016). Naive Human Pluripotent Cells Feature a Methylation Landscape Devoid of Blastocyst or Germline Memory. Cell Stem Cell 18, 323–9.

Patel, S., Bonora, G., Sahakyan, A., Kim, R., Chronis, C., Langerman, J., Fitz-Gibbon, S., Rubbi, L., Skelton, R. J., Ardehali, R. et al. (2017). Human Embryonic Stem Cells Do Not Change Their X Inactivation Status during Differentiation. Cell Rep 18, 54–67.

Petropoulos, S., Edsgard, D., Reinius, B., Deng, Q., Panula, S. P., Codeluppi, S., Plaza Reyes, A., Linnarsson, S., Sandberg, R. and Lanner, F. (2016). Single-Cell RNA-Seq Reveals Lineage and X Chromosome Dynamics in Human Preimplantation Embryos. Cell 165, 1012–26.

Quenneville, S., Verde, G., Corsinotti, A., Kapopoulou, A., Jakobsson, J., Offner, S., Baglivo, I., Pedone, P. V., Grimaldi, G., Riccio, A. et al. (2011). In embryonic stem cells, ZFP57/KAP1 recognize a methylated hexanucleotide to affect chromatin and DNA methylation of imprinting control regions. Mol Cell 44, 361–72.

Reik, W. and Kelsey, G. (2014). Epigenetics: Cellular memory erased in human embryos. Nature 511, 540–541.

Roode, M., Blair, K., Snell, P., Elder, K., Marchant, S., Smith, A. and Nichols, J. (2012). Human hypoblast formation is not dependent on FGF signalling. Dev Biol 361, 358–63.

Rossant, J. and Tam, P. P. L. (2017). New Insights into Early Human Development: Lessons for Stem Cell Derivation and Differentiation. Cell Stem Cell 20, 18–28.

Sahakyan, A., Kim, R., Chronis, C., Sabri, S., Bonora, G., Theunissen, T. W., Kuoy, E., Langerman, J., Clark, A. T., Jaenisch, R. et al. (2017). Human Naive Pluripotent Stem Cells Model X Chromosome Dampening and X Inactivation. Cell Stem Cell 20, 87–101.

Semrau, S., Goldmann, J., Soumillon, M., Mikkelsen, T. S., Jaenisch, R. and van Oudenaarden, A. (2016). Dynamics of lineage commitment revealed by single-cell transcriptomics of differentiating embryonic stem cells. bioRxiv.

Sherry, S. T., Ward, M. H., Kholodov, M., Baker, J., Phan, L., Smigielski, E. M. and Sirotkin, K. (2001). dbSNP: the NCBI database of genetic variation. Nucleic Acids Res 29, 308–11.

Silva, S. S., Rowntree, R. K., Mekhoubad, S. and Lee, J. T. (2008). X-chromosome inactivation and epigenetic fluidity in human embryonic stem cells. Proc Natl Acad Sci U S A 105, 4820–5.

Smallwood, S. A., Lee, H. J., Angermueller, C., Krueger, F., Saadeh, H., Peat, J., Andrews, S. R., Stegle, O., Reik, W. and Kelsey, G. (2014). Single-cell genome-wide bisulfite sequencing for assessing epigenetic heterogeneity. Nat Methods 11, 817–20.

Smith, A. (2017). Formative pluripotency: the executive phase in a developmental continuum. Development 144, 365–373.

Sperber, H., Mathieu, J., Wang, Y., Ferreccio, A., Hesson, J., Xu, Z., Fischer, K. A., Devi, A., Detraux, D., Gu, H. et al. (2015). The metabolome regulates the epigenetic landscape during naive-to-primed human embryonic stem cell transition. Nat Cell Biol 17, 1523–1535.

Takahashi, K., Tanabe, K., Ohnuki, M., Narita, M., Ichisaka, T., Tomoda, K. and Yamanaka, S. (2007). Induction of pluripotent stem cells from adult human fibroblasts by defined factors. Cell 131, 861–72.

Takashima, Y., Guo, G., Loos, R., Nichols, J., Ficz, G., Krueger, F., Oxley, D., Santos, F., Clarke, J., Mansfield, W. et al. (2014). Resetting Transcription Factor Control Circuitry toward Ground-State Pluripotency in Human. Cell 158, 1254–1269.

Tang, W. W., Dietmann, S., Irie, N., Leitch, H. G., Floros, V. I., Bradshaw, C. R., Hackett, J. A., Chinnery, P. F. and Surani, M. A. (2015). A Unique Gene Regulatory Network Resets the Human Germline Epigenome for Development. Cell 161, 1453–67.

Tesar, P. J., Chenoweth, J. G., Brook, F. A., Davies, T. J., Evans, E. P., Mack, D. L., Gardner, R. L. and McKay, R. D. (2007). New cell lines from mouse epiblast share defining features with human embryonic stem cells. Nature 448, 196–9.

Theunissen, T. W., Friedli, M., He, Y., Planet, E., O’Neil, R. C., Markoulaki, S., Pontis, J., Wang, H., Iouranova, A., Imbeault, M. et al. (2016). Molecular Criteria for Defining the Naive Human Pluripotent State. Cell Stem Cell 19, 502–515.

Theunissen, T. W., Powell, B. E., Wang, H., Mitalipova, M., Faddah, D. A., Reddy, J., Fan, Z. P., Maetzel, D., Ganz, K., Shi, L. et al. (2014). Systematic identification of culture conditions for induction and maintenance of naive human pluripotency. Cell Stem Cell 15, 471–87.

Thomson, J. A., Itskovitz-Eldor, J., Shapiro, S. S., Waknitz, M. A., Swiergiel, J. J., Marshall, V. S. and Jones, J. M. (1998). Embryonic stem cell lines derived from human blastocysts. Science 282, 1145–7.

Vallot, C., Patrat, C., Collier, A. J., Huret, C., Casanova, M., Liyakat Ali, T. M., Tosolini, M., Frydman, N., Heard, E., Rugg-Gunn, P. J. et al. (2017). XACT Noncoding RNA Competes with XIST in the Control of X Chromosome Activity during Human Early Development. Cell Stem Cell 20, 102–111.

Van Der Maaten, L. (2014). Accelerating t-SNE using tree-based algorithms. Journal of Machine Learning Research 15, 3221–3245.

van der Maaten, L. and Hinton, G. (2008). Visualizing data using t-SNE. Journal of Machine Learning Research 9, 2579–2605.

von Meyenn, F., Berrens, R. V., Andrews, S., Santos, F., Collier, A. J., Krueger, F., Osorno, R., Dean, W., Rugg-Gunn, P. J. and Reik, W. (2016). Comparative Principles of DNA Methylation Reprogramming during Human and Mouse In Vitro Primordial Germ Cell Specification. Dev Cell 39, 104–115.

Walter, M., Teissandier, A., Perez-Palacios, R. and Bourc’his, D. (2016). An epigenetic switch ensures transposon repression upon dynamic loss of DNA methylation in embryonic stem cells. Elife 5.

Wang, J., Xie, G., Singh, M., Ghanbarian, A. T., Rasko, T., Szvetnik, A., Cai, H., Besser, D., Prigione, A., Fuchs, N. V. et al. (2014). Primate-specific endogenous retrovirus-driven transcription defines naive-like stem cells. Nature 516, 405–9.

Ware, C. B., Wang, L., Mecham, B. H., Shen, L., Nelson, A. M., Bar, M., Lamba, D. A., Dauphin, D. S., Buckingham, B., Askari, B. et al. (2009). Histone deacetylase inhibition elicits an evolutionarily conserved self-renewal program in embryonic stem cells. Cell Stem Cell 4, 359–69.

Wray, J., Kalkan, T. and Smith, A. G. (2010). The ground state of pluripotency. Biochem Soc Trans 38, 1027–32.

Wu, J., Okamura, D., Li, M., Suzuki, K., Luo, C., Ma, L., He, Y., Li, Z., Benner, C., Tamura, I. et al. (2015). An alternative pluripotent state confers interspecies chimaeric competency. Nature 521, 316–321.

Yan, L., Yang, M., Guo, H., Yang, L., Wu, J., Li, R., Liu, P., Lian, Y., Zheng, X., Yan, J. et al. (2013). Single-cell RNA-Seq profiling of human preimplantation embryos and embryonic stem cells. Nat Struct Mol Biol 20, 1131–9.

Yang, Y., Liu, B., Xu, J., Wang, J., Wu, J., Shi, C., Xu, Y., Dong, J., Wang, C., Lai, W. et al. (2017). Derivation of Pluripotent Stem Cells with In Vivo Embryonic and Extraembryonic Potency. Cell, 169 243–257 e25.

Ying, Q. L., Wray, J., Nichols, J., Batlle-Morera, L., Doble, B., Woodgett, J., Cohen, P. and Smith, A. (2008). The ground state of embryonic stem cell self-renewal. Nature 453, 519–23.

Yu, J., Vodyanik, M. A., Smuga-Otto, K., Antosiewicz-Bourget, J., Frane, J. L., Tian, S., Nie, J., Jonsdottir, G. A., Ruotti, V., Stewart, R. et al. (2007). Induced pluripotent stem cell lines derived from human somatic cells. Science 318, 1917–20.

Zhou, W., Choi, M., Margineantu, D., Margaretha, L., Hesson, J., Cavanaugh, C., Blau, C. A., Horwitz, M. S., Hockenbery, D., Ware, C. et al. (2012). HIF1alpha induced switch from bivalent to exclusively glycolytic metabolism during ESC-to-EpiSC/hESC transition. EMBO J 31, 2103–16.

Zimmerlin, L., Park, T. S., Huo, J. S., Verma, K., Pather, S. R., Talbot, C. C., Agarwal, J., Steppan, D., Zhang, Y. W., Considine, M. et al. (2016). Tankyrase inhibition promotes a stable human naïve pluripotent state with improved functionality. Development 143, 4368–4380.

